# Highly Connected and Highly Variable: A Core Brain Network during Resting State Supports Propofol-induced Unconsciousness

**DOI:** 10.1101/2022.03.03.482914

**Authors:** Siyang Li, Yali Chen, Peng Ren, Zhipeng Li, Jun Zhang, Xia Liang

**Affiliations:** School of Life Science and Technology, Harbin Institute of Technology, Harbin, China; Institute of Space Environment and Materiel Science, Harbin Institute of Technology, Harbin, China; Department of Anesthesiology, Shanghai Cancer Center, Fudan University, Shanghai, China; Department of Oncology, Shanghai Medical College, Fudan University, Shanghai, China

**Keywords:** Consciousness, neural correlates of consciousness, rich-club, modular variability, dynamic states

## Abstract

Leading theories of consciousness make diverging predictions for where and how neural activity gives rise to subjective experience. The Global Neuronal Workspace theory (GNW) states that consciousness is instantiated through global broadcasting of information across the prefrontal-parietal regions, whereas the integrated information theory (IIT) postulates that consciousness requires the posterior cortex to produce maximally irreducible integrated information. As both theories seem to partially agree on that the neural correlates of consciousness (NCC) require globally integrated brain activity across a network of functionally specialized modules, it is not known yet whether brain regions with such functional configurations would align with the NCC distribution predicted by the GNW or the IIT. We scanned resting-state fMRI data from 21 subjects during wakefulness, propofol-induced sedation and anesthesia. Graph-theoretical analysis were conducted on awake fMRI data to search for the NCC candidates as brain regions that exhibit both high rich-clubness and high modular variability. Another independent dataset of 10 highly-sampled subjects were used to validate the NCC distribution at individual-level. Brain module-based dynamic analysis was conducted to estimate temporal stability of the NCC candidates. Alterations in functional connectivity and modular variability from awake to propofol-induced anesthesia were assessed to test the involvement of the NCC candidates in conscious processing. NCC candidates that are characterized by both high functional interconnectivity and high modular variability were identified to locate in prefrontal and temporoparietal cortices, which covered brain structures predicted by the GNW as well as the IIT. The identified NCC was found to mainly attributed to higher-order cognitive functions, and associated with genes enriched in synaptic transmission. Dynamic analysis revealed two discrete reoccurring brain states, which were characterized by their difference in temporal stability — the state dominated by the NCC candidates appearred to be temporally more stable than the other state predominately composed of primary sensory/motor regions, suggesting that the identified NCC members could sustain conscious contents as metastable network representations. Finally, we showed that the prefrontal GNW regions and posterior IIT regions within the identified NCC was differentially modulated in terms of functional connectedness and modular variability in response to loss of consciousness induced by propofol anesthesia. This work offers a framework to search for neural correlates of consciousness by charting the brain network topology, and provides new insights in understanding the distinct roles of the frontoparietal and posterior network in underpinning human consciousness.

**Highlights:** Studies suggest that there are neural correlates of consciousness (NCC) we experience subjectively everyday. By overlapping regions with both high functional interconnectivity (rich-clubness) and high modular variability, we identified the putative NCC distributed in prefrontal and temporoparietal cortices, attributed to higher-order cognitive functions, and associated with genes enriched in synaptic transmission. We further revealed that the NCC members appeared to sustain conscious contents as metastable network representations in a reoccurring NCC dominant state. The identified NCC architecture was significantly modulated in terms of functional connectedness and modular varibility during propofol anesthesia, demonstrating its critical role in supporting consciousness. These findings testify to the NCC’s abilities in information integration and differentiation, and provide novel insights in reconciling the ongoing discussion of the contribution of anterior versus posterior regions in supporting human consciousness.

## Introduction

Consciousness is what we experience subjectively every day when a salient external stimulus accesses conscious operation, or when our minds spontaneously wander into episodic memory. While mounting evidence suggests the existence of the neural correlates of consciousness (NCC), it is not known yet where specially they are located in the brain ^1^. Different theories of consciousness make diverging predictions for where and how neural activity gives rise to subjective experience ^2-5^. The Global Neuronal Workspace theory (GNWT) states that consciousness is instantiated through global broadcasting of information across a core set of highly interconnected areas, including predominately the prefrontal-parietal regions^6-8^. In contrast, the integrated information theory (IIT) postulates that consciousness requires both integration and differentiation to produce maximally irreducible integrated information generated by the posterior hot zone of the temporo-parieto-occipital areas ^9^.

Despite various distinctions among these influential theories of consciousness, they seem to partially agree on an account that the neural correlates of consciousness require to support globally integrated brain activity across a network of functionally specialized modules^6,7,10-12^. Indeed, prior studies estimating brain integration based on graph-theoretical tools have confirmed the reduction in brain-wide global sychronization under diverse general anesthetics and in patients with disorders of consciousness^6,7,13-20^. In contrast, activities of functional separated modules, another important aspect involving in the information integration process during conscious experience, have received little attention in previous studies searching for the neural correlates of consciousness. The brain’s modular structure represents groups of densely connected regions with shared functionality ^21-24^, and has become the focus of many investigations. By demonstrating remarkable temporal and individual variability in the spatial topography of brain modules ^25-30^, recent studies have suggested that variability in modular topography reflects flexible adaptation to changing environment and underlies individual differences in cognitive and behavioral capacity ^27,30^. However, the relevance of a region’s modular variability to its role in consciousness remains an open question. Given the evidence that regions with higher modular variability are more likely to be reassigned depending on the mental context, consciously accessible experiences fluctuating over time or among individuals would require diversified representations supported by diverse module patterns. We therefore hypothesize that, apart from the centralized connectivity architecture, neural correlates of consciousness should also be characterized by higher variability of modular affiliations in response to various external stimulus, or to stream of spontaneous thoughts during wakeful resting state.

To test our hypothesis, we aimed to locate neural correlates of consciousness by analyzing two critical organizational aspects, i.e., functional integration and diversification, of spontaneous brain activity from resting-state fMRI data. We searched for NCC candidates by identifying brain regions with both high rich-clubness and high modular variability in a cohort of 21 subjects during awake resting-state, and validated at individual-level in another independent dataset of 10 highly-sampled subjects. We also explored the dynamics of spontaneous brain networks, and hypothesized that the identified NCC candidates should be more involved in a temporally more stable state. To further demonstrate the linkage between NCC candidates and conscious processing, we analyzed fMRI data during propofol administration with varied concentrations, and hypothesized that the NCC regions would be more sensitive to general anesthetic-induced loss of consciousness.

## Results

### Identification of neural correlates of consciousness

To identify brain regions that may act as NCC, we explored two fundamental topological properties of functional brain networks: the rich-club structure and the modular variability. NCC candidates are identified as brain regions that (a) belong to the “rich-club” structure, and (b) exhibit high variability in modular topography across both individuals and time (Figure 1).

**Figure 1.**
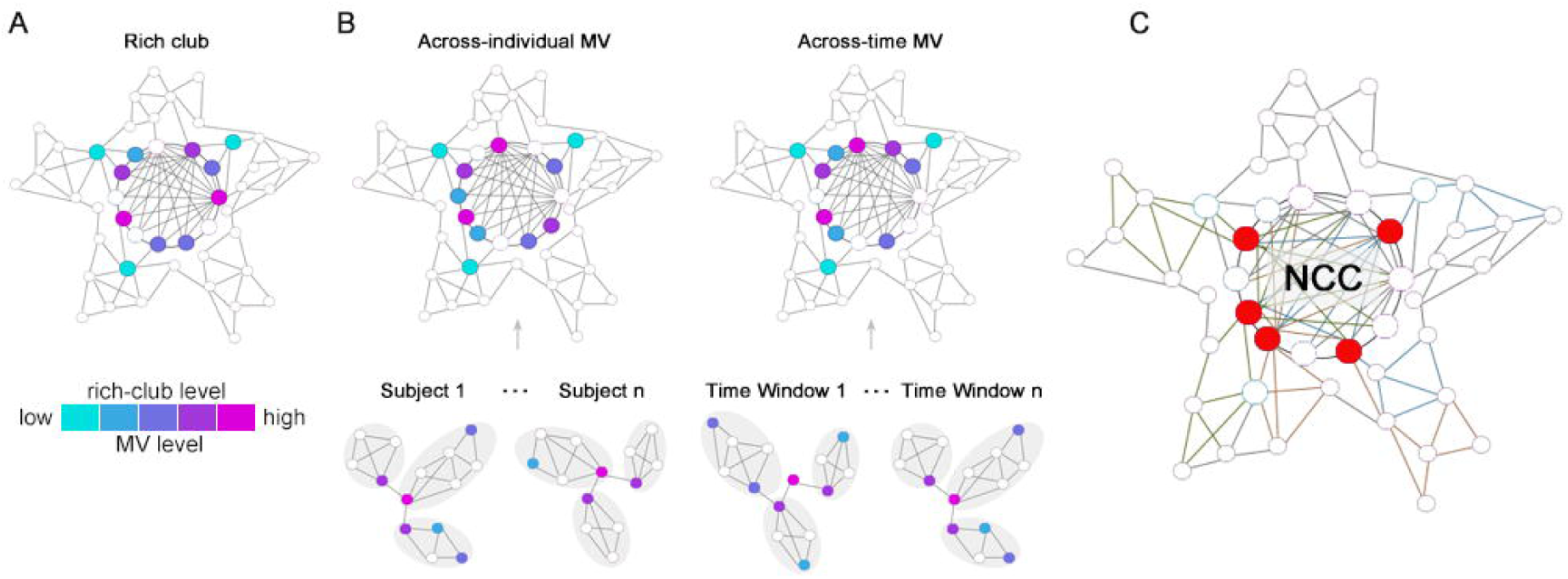
Schematic of the identification of neural correlates of consciousness (NCC). **(A)** Rich-club consists of high-degree hub regions that are more densely interconnected among themselves. **(B)** Brain regions that tend to sway between different modules across different subjects or different time windows are characterized by high across-individual or across-time modular variability. **(C)** Regions with high rich-club level, high across-individual MV as well as high across-time MV were overlapped to give rise to a putative NCC template.

The rich-club organization and modular variability were analyzed on the group-averaged functional brain networks in twenty-one subjects during awake. Consistent with previous studies, we demonstrated that the functional brain networks exhibited significant rich-club structure comparing to randomly reorganized networks for a range of connectivity strength *s* = 1 to *s* = 30 [*p* < 0.001, permutation test (*see Materials and Methods*), Figure S1]. By showing the anatomical distribution of rich-club members at different levels (top 10% to 50% connectivity strength, Figure 2A), we observed a tendency of hierarchical layout for rich-club areas. While the primary sensorimotor, auditory regions are among the lower level of connectivity strength (top 40% ∼ 50%), followed by association cortices of dorsolateral frontoparietal regions (top 20% ∼ 40%), the most highly-connected rich-club core (≥top 20%) is occupied by the anterior and posterior midline, bilateral temporoparietal junction areas, forming the default mode network (DMN).

**Figure 2.**
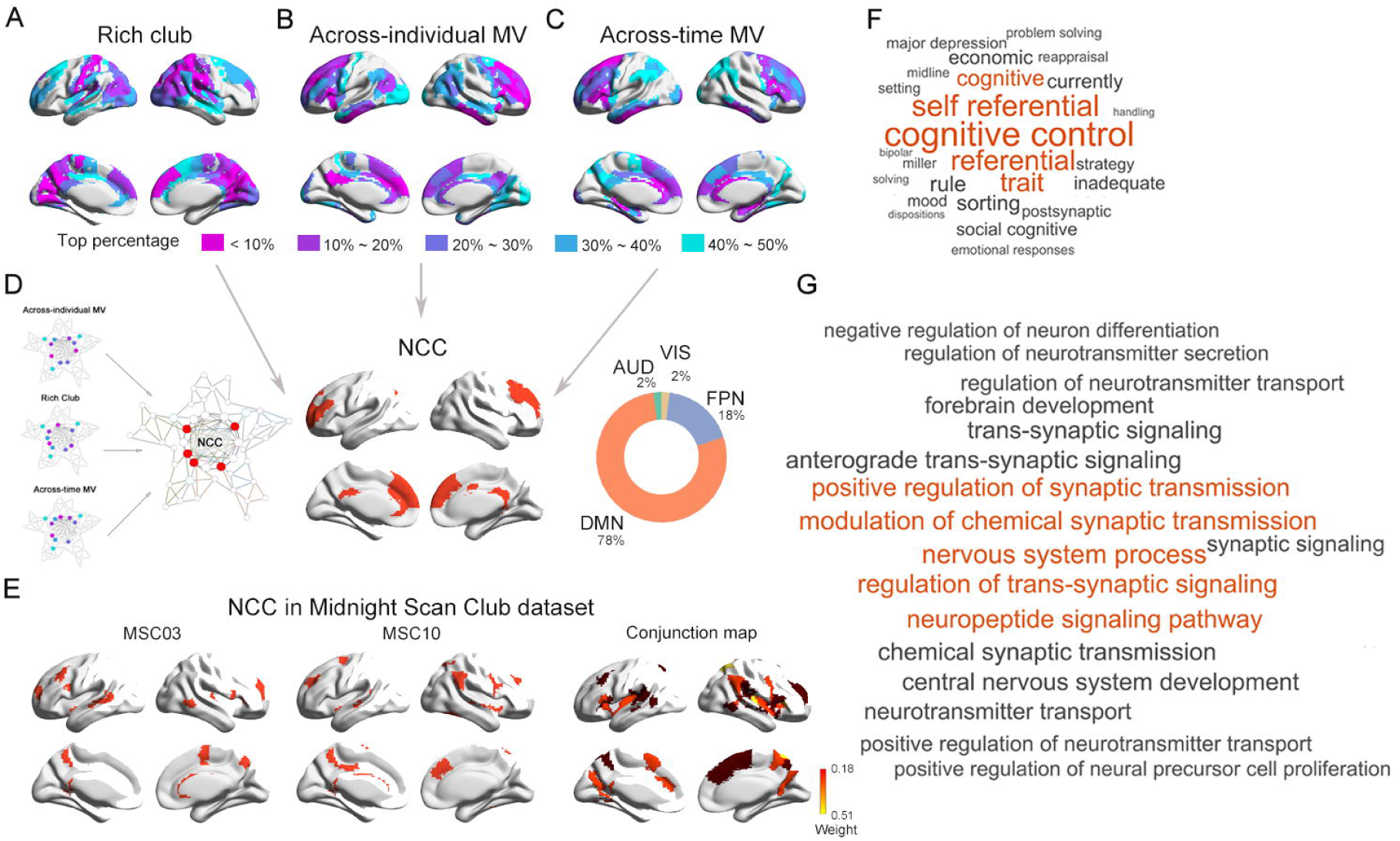
Neural correlates of consciousness (NCC) identification and functional decoding. **(A-C)** Anatomical distribution of rich-club, across-individual MV and across-time MV members at different levels (top 10% to 50%). **(D)** A putative NCC template by overlapping the three maps of regions with high rich-club connectedness (top 40%), high across-individual MV (top 40%) and high across-time MV (top 40%), 78% of which was in DMN and 18% was in FPN. **(E)** Individual-level GNW map for an example subject (MSC03) (upper) and the conjunction map of NCC map across individuals (lower) from the MSC dataset. **(F)** Cognitive terms that are closely related to the putative NCC regions. Terms that are of higher magnitude of the similarity with the NCC distribution were displayed with larger font size. **(G)** Gene-ontology enrichment results for the differentially expressed transcriptional profiles between NCC and non-NCC regions. Neural processes that are more significantly enriched in GO analysis were displayed with larger font size.

We next explored modular organization of the group-averaged functional brain networks. We detected seven modules during awake resting-state (Figure S2), including the frontoparietal network (FPN), default mode network (DMN), auditory network (AUD), sensorimotor network (SMN), visual network (VIS), medial temporal lobe (MTL) and subcortical network. Modular structure was also evaluated for each subject and modular variability (MV) was computed separately across individuals and time (*see Materials and Methods*). Figure 2B and C showed the anatomical layout of brain regions with different level of across-individual and across-time MV (top 10% to 50%). Regions in the frontal, temporoparietal and insular cortices exhibit high variability in modular affiliation (≥top 20%), especially the medial and lateral prefrontal areas, are among the highest in across-individual and across-time MV (top 10%).

We then overlapped the above three maps of regions with high rich-club connectedness (top 40%), high across-individual MV (top 40%) and high across-time MV (top 40%), which gave rise to a putative NCC template of twenty brain areas (8% of all brain regions) distributing primarily in the anterior and posterior midline, lateral prefrontal and temporoparietal junction areas (Figure 2D). Remarkably, a large percent of these identified regions resided within the DMN (78%) and FPN (18%), which covered most brain structures matching the GNW as well as the ITT prediction. We also generated overlapping maps of the three topological metrics using different percentile (top 30% and top 50%) of regions with top rich-clubness and MV (Figure S3), and validated the robustness of the putative NCC candidates, especially the prefrontal regions that were consistently detected as potential NCC with both high interconnectivity and high modular variability.

To further validate our findings and to explore individual differences in NCC constitution, we used the second, independent dataset of ten highly-sampled individuals scanned for five hours (Midnight Scan Club dataset, *see Materials and Methods*) to identify individual-level NCC with both high rich-club connectedness (top 40%) and high MV (top 40%). Although there were variations across the individual-level NCC map (Figure S4), their conjunction map showed that the candidate NCC regions in more than 30% of subjects were predominately distributed in anterior, middle and posterior cingulate cortices, middle frontal and temporoparietal cortices, and medial occipital gyrus (Figure 2E), which largely resembles the NCC distribution observed in our main dataset.

### Functional decoding of the NCC candidates

To understand cognitive functions served by NCC, we conducted a term-based meta-analysis using NeuroQuery database ^31^. By examining the association between a large repertoire of cognitive and neurological terms with the putative NCC template created from the above step, we revealed that our identified NCC regions are most closely related to higher cognitive functions such as ‘cognitive control’, ‘self-referential’ and ‘trait’ (Figure 2F). Meta-analysis was also performed using the NeuroSynth database ^32^, which yielded similar functional decoding result (Figure S5).

### Transcriptional origins of the NCC candidates

To understand genetic basis underlying the NCC structure, we explored differential gene expression between the NCC and non-NCC regions using regional expression profiles of 15655 genes from Allen Human Brain Atlas (AHBA) dataset ^33^. Our results revealed that, comparing with the non-NCC regions, genes in the NCC regions were enriched in neuron transmission and signaling functions (Figure 2G), such as ‘regulation of trans-synaptic signaling’ and ‘modulation of chemical synaptic transmission’ (*P* < 0.05).

### NCC candidates are more involved in a dynamically metastable state

In sought to explore the dynamics of conscious processing, we developed an approach to identify “brain states” by examining time-varying reconfigurations in brain modular structure variability (*see Material and Methods for details*). The “brain states” represented elementary functional configurations of the brain, in which time windows with similar temporal variability (i.e., similar stability) in modular affiliations were clustered together. We identified five brain states at the group-level. When projecting each brain state back to individuals, three states were presented in very few subjects, and were thus discarded, resulting in two dominant brain states retained for further analysis. For each state, we evaluated their temporal stability, calculated as the proportion of time windows showing the least MV across time (*see Material and Methods*). We found that one of the two dominant brain states (state 1) exhibited higher stability than the other (state 2; Figure 3A left; *t* = 12.05, *P* < 10^−5^), suggesting that brain state 1 may represent a temporally more stable state.

**Figure 3.**
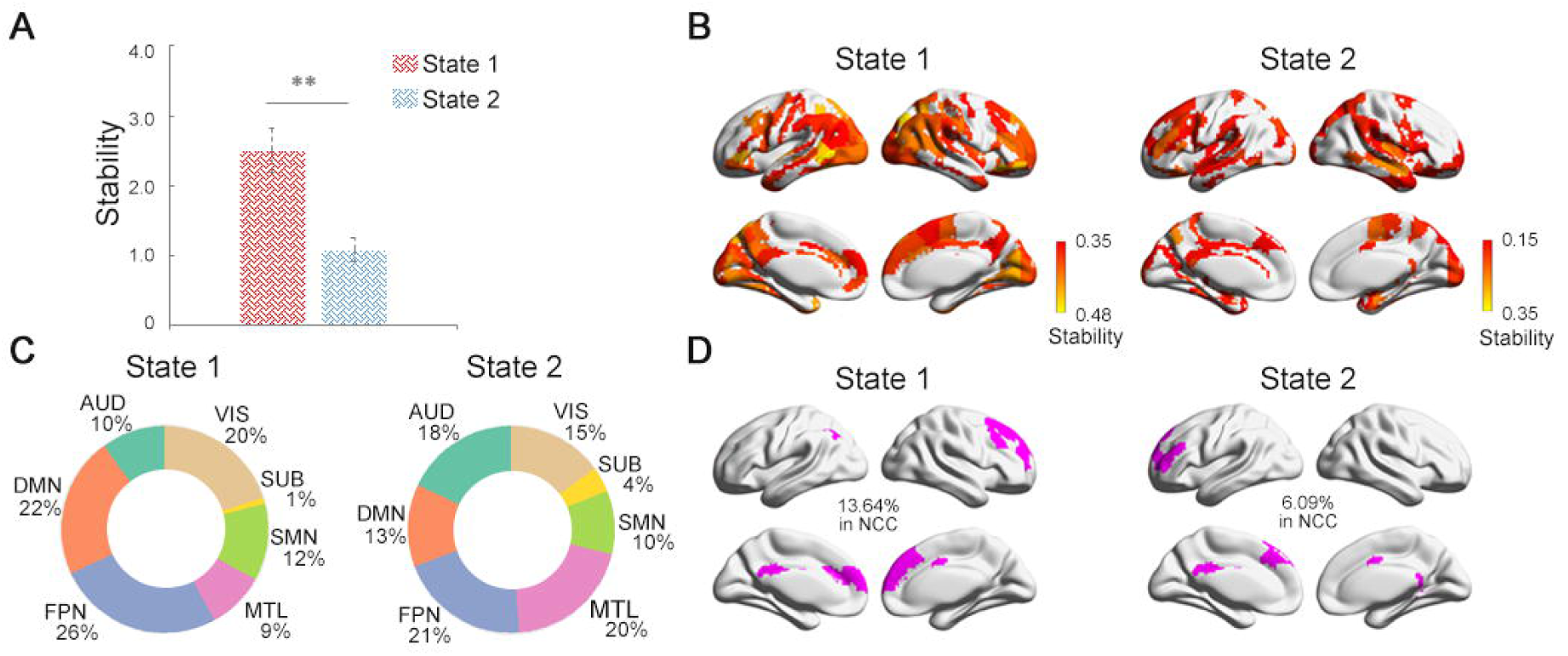
Neural correlates of consciousness (NCC) dynamics. **(A)** Two different iterating activity states with state 1 having higher stability than state 2. **(B)** Brain regions with high stability (>mean) and **(C)** their percentile distribution across brain modules in state 1 (left) and state 2 (right). **(D)** Overlaps between identified NCC space with the stable regions in state 1 (left) and state 2 (right).

Figure 3B showed the brain regions with high stability (> mean) in each state. In state 1, we found that the most stable brain regions are preferentially distributed in the DMN and FPN modules, taking about 48% of all the stable regions (Figure 3C). In contrast, in state 2, the most stable regions were dominated by primary sensory and limbic networks (Figure 3B), with the DMN and FPN modules taking a less proportion (34% in total; Figure 3C right). By overlapping our identified NCC candidates with the stable regions in each state (Figure 3D), we observed that while both states exhibit significant overlaps with the NCC in regions of prefrontal cortex, anterior and posterior cingulate cortices (*Ps* < 0.001, permutation test), the stable regions in state 1 shared 13.64% with the NCC, which was significantly more (*P* < 0.001, permutation test) than that in state 2 (6.09%). We also evaluated the overlap between stable regions in each state and NCC candidates identified at other rich-club and MV levels, and validated that the NCC overlapped more with stable regions in state 1 than in state 2 (Figure S6). Taking together, our results demonstrated that the spontaneous brain activities could be decomposed into a more stable brain state, steered predominately by the NCC members, and another less stable state dominated by primary sensorimotor regions which may be associated with subliminal processing.

### NCC signatures track loss of consciousness during propofol administration

Having identified the NCC and linked it with the metastable dynamic brain state, we next sought to investigate the role of the NCC in supporting human consciousness by comparing fMRI data in the same cohort of subjects from awake to sedation and unconscious state induced by different concentrations of intravenous anesthetic propofol. The identification of NCC allowed for the classification of brain regions as NCC and non-NCC members, and connections into 3 categories, including NCC connections linking NCC members, feeder connections linking NCC and non-NCC regions, and local connections linking non-NCC regions. We therefore calculated MV for NCC and non-NCC regions, and FC strength (FCS) for the 3 categories of connections, respectively, and expected to observe alterations, especially within the NCC, during anesthesia-induced loss of consciousness.

One-way ANOVA revealed marginal significant effect of anesthesia in static FC strength (Figure 4A) of local connections (*F (2,26) =* 3.59, *P*_*uncorrected*_ *=* 0.04, *P*_*corrected*_ *=* 0.12), and marginal significant effect of anesthesia in NCC connections (*F (2,26) =* 3.57, *P*_*uncorrected*_ *=* 0.04, *P*_*corrected*_ *=* 0.12), both of which decreased with loss of consciousness. Post-hoc tests found no significant changes in NCC or local connections between each pair of the three conscious levels. To further explore which specific connections are impacted during propofol administration, we performed network-based statistics (NBS) between different conscious levels for NCC and local connections, respectively. Our results for NCC connections revealed significant decreases in strength of FC among the midline key regions of the DMN, including the medial prefrontal cortex (mPFC), anterior cingulate cortex (ACC) and posterior cingulate cortex (PCC), between wakefulness and anesthesia (*P* < 0.005), and among the midline DMN areas and lateral middle frontal gyrus (MFG) between sedation and anesthesia (*P* < 0.005, Figure 4C). As for the local connections (Figure S7), reductions in FC were mainly distributed among SMN, VIS and MTL networks (*P* < 0.005).

**Figure 4.**
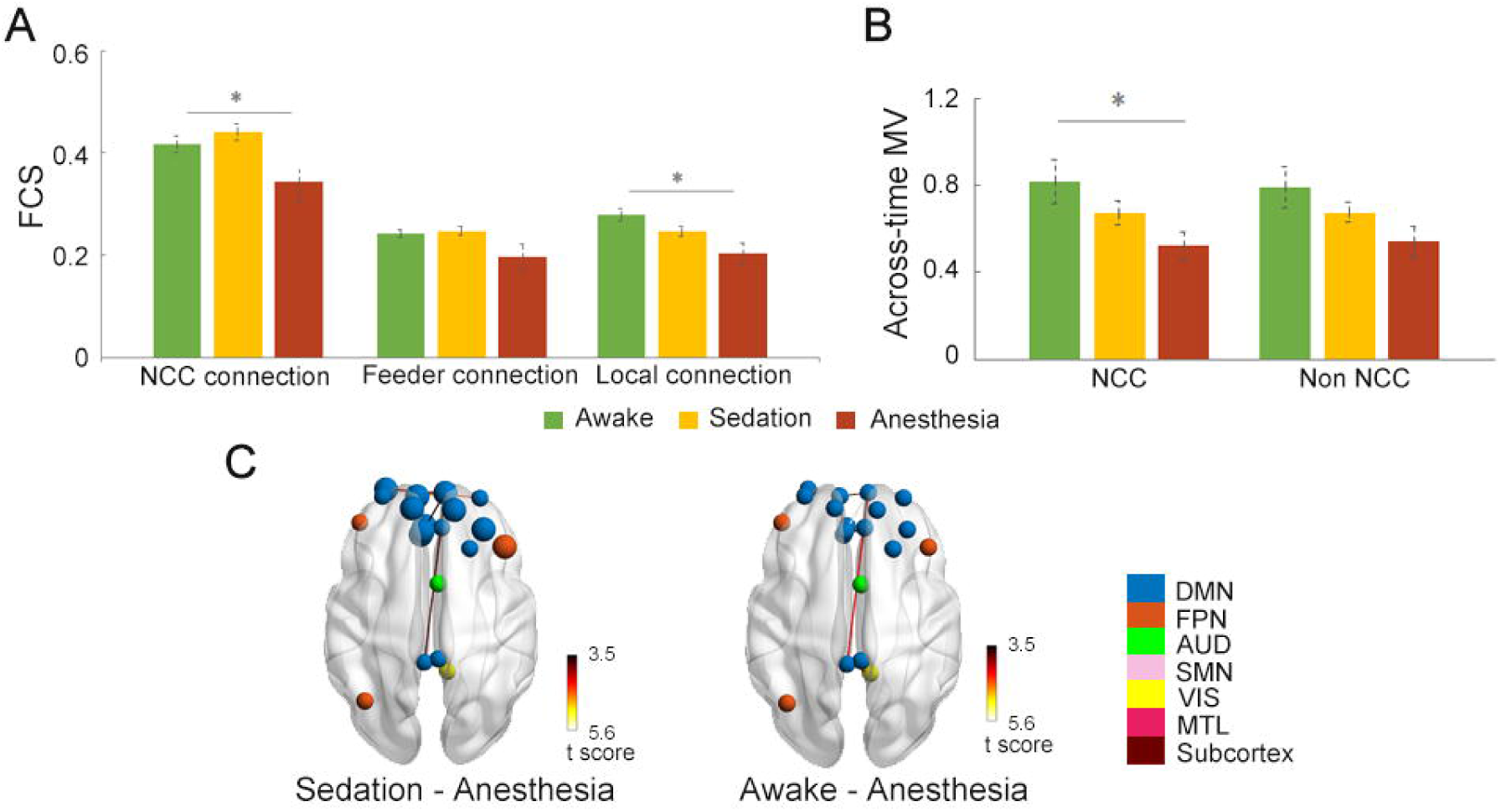
Static NCC signatures track loss of consciousness during Propofol anesthesia. **(A)** Differences in static FC of NCC, feeder, and local connections from awake to sedation and unconsciousness. **(B)** Differences in across-time MV of NCC and non-NCC regions from awake to sedation and unconsciousness. **(C)** NCC connections showing significant reductions in NBS between awake and unconsciousness, and NCC regions with significant decreases in across-time MV from sedation to anesthesia condition (highlighted in larger circles). Color of nodes indicates modular affiliation. *\*,P*_*uncorrected*_ *< 0*.*05*.

We also observed marginal significant effect of loss of consciousness in across-time MV only within the NCC (*F (2,26) =* 3.76, *P*_*uncorrected*_ *=* 0.037, *P*_*corrected*_ *=* 0.074), with the across-time MV decreased as the dose of anesthesia deepened (Figure 4B). Post-hoc tests found no significant changes between pairs of the three conscious levels. Further regional-wise paired t comparisons revealed significant reductions of across-time MV locating primarily in the medial and lateral prefrontal areas from sedation to anesthesia (*P*_*corrected*_ *< 0*.*05*, Figure 4C), indicating the evident degeneration of the ability for these regions to dynamically vary its modular affiliation across time. No significant effect of consciousness was found for across-individual MV.

Dynamic analysis of brain states revealed that significant anesthesia-related changes in FC strength were observed only during the stable state (state 1) within the NCC (*F (2,23) =* 7.59, *P*_*uncorrected*_ *=* 0.003, *P*_*corrected*_ *=* 0.009, Figure 5A). Marginal significant effect of anesthesia was also observed for feeder (*F (2,23) =* 3.84, *P*_*uncorrected*_ *=* 0.038, *P*_*corrected*_ *=* 0.11) and local connections (*F (2,23) =* 3.77, *P*_*uncorrected*_ *=* 0.040, *P*_*corrected*_ *=* 0.12) during the stable state. Post-hoc tests revealed significant decreases in NCC connection from sedation to anesthesia (*t* = 4.37, *P*_*uncorrected*_ *=*0.001, *P*_*corrected*_ = 0.003) during the stable state, while no significant between-condition differences were observed for feeder or local connections. Further NBS analysis for NCC connections in state 1 revealed significant decreases among regions of mPFC, ACC, PCC and MFG (*P* < 0.005, Figure 5B). As for the feeder connections, significant reductions were mostly between NCC regions with non-NCC regions in DMN from awake to anesthesia, and with non-NCC regions in FPN from sedation to anesthesia (*P* < 0.005, Figure 5B). Local connections among a cluster of regions across the posterior part of the brain were shown to reduce from awake and sedation to anesthesia (*P* < 0.005, Figure S7). No significant effect of anesthesia was observed in across-time MV in either of the two dynamic states. Similar anesthesia effects were also observed using the NCC identified at different rich-club and MV levels (Figure S8).

**Figure 5.**
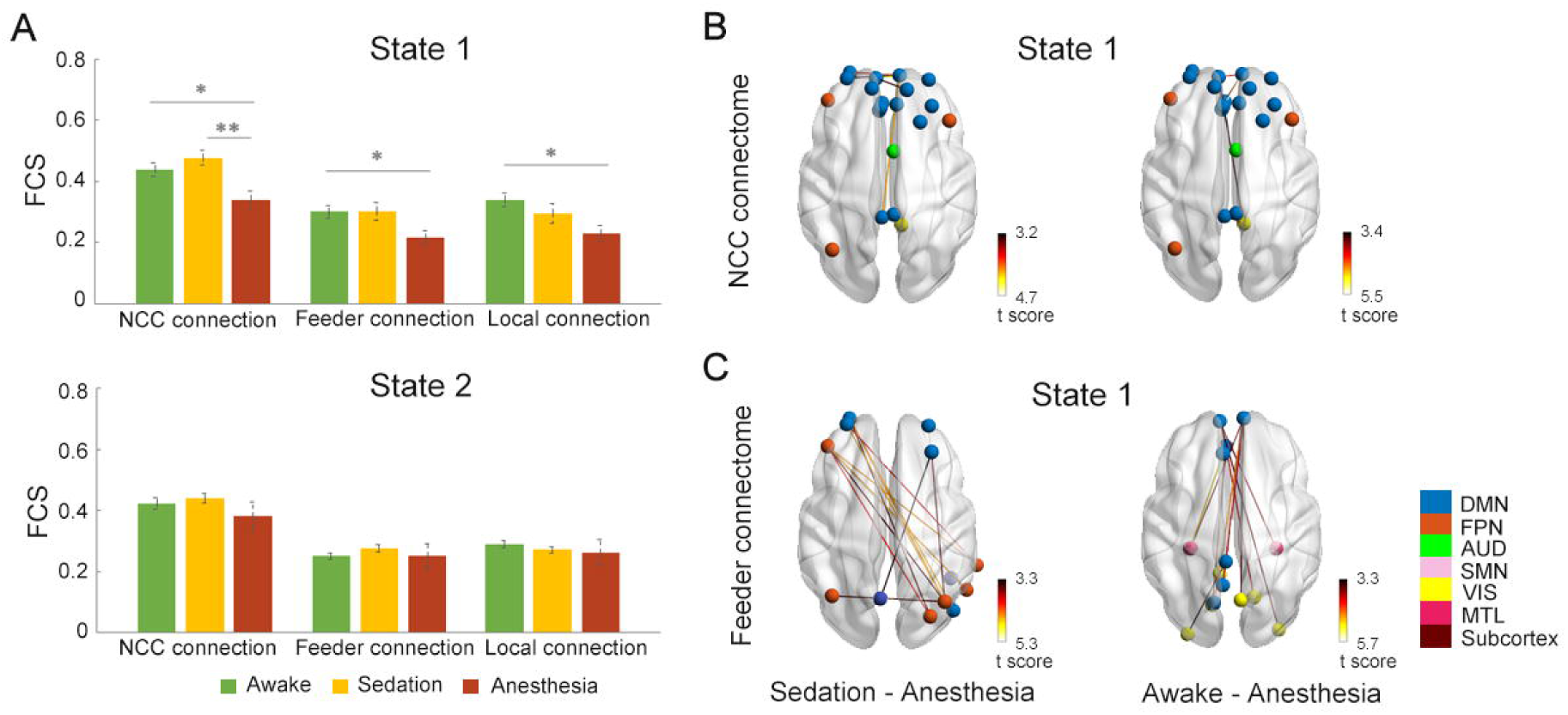
Dynamic NCC signatures track loss of consciousness during Propofol anesthesia. **(A)** Differences in dynamic FC of NCC, feeder, and local connections from awake to sedation and unconsciousness during each state. **(B)** NCC connections showing significant reductions in NBS between different conscious levels during state 1. **(C)** Feeder connections showing significant reductions in NBS between different conscious levels during state 1. Color of nodes indicates modular affiliation. *\*, P*_*uncorrected*_ *< 0*.*05; **, P*_*corrected*_ *< 0*.*05*.

## Discussion

The present study proposes to locate the neural correlates of consciousness by taking advantage of recent developments in network neuroscience. We identified a cluster of regions distributed in prefrontal and temporoparietal cortices that are characterized by both high functional interconnectivity and high modular variability to constitute the putative NCC. Dynamic analysis revealed two discrete reoccurring brain states, which are characterized by their differences in temporal stability — the state dominated by the identified NCC appears to be temporally more stable than the other state predominately composed of primary regions, demonstrating that the identified neural correlates of consciousness are able to sustain conscious contents as metastable network representations. We also demonstrated that the identifed NCC members are associated with overexpressed genes enriched in synaptic transmission, supporting its role in higher-order cognitive functions revealed in task-based meta-analysis. Finally, we showed that the identified NCC candidates were significantly modulated in terms of functional connectedness and modular variability in response to loss of consciousness during propofol anesthesia.

Combing two well-defined graph-theoretical network measures, we located the putative NCC members in a distributed set of regions residing predominately in the default-mode and frontoparietal networks. While the identified regions included the posterior cingulate and temporoparietal cortices, which are main candidates of the “posterior hot zone” advocated by the IIT, the inclusion of anterior areas in the putative NCC provides support for the GNW theory. Notably, regions in the anterior cortex were observed to show both the densest connectivity and the highest modular variability, situating at the core of the putative NCC. This is in well concordance with initial theoretical and simulation studies of GNW predicting the dorsolateral prefrontal and anterior cingulate cortices to be the major contributors of the workspace ^34^, and is also consistent with recent efforts in quantifying the global workspace ^35,36^ by considering the brain as an information processing system. The posterior cortical regions, on the other hand, were of high connectivity but relatively low temporal variability in modular structure. This is in line with a recent study^36^ demonstrating that the default mode system, especially its posterior regions, exhibited higher prevalence of synergistic over redundant information, which suggests an integrating rather than broadcasting role of these regions in processing information accessible to consciousness. Thus, the PCC may act as an integrator with rich connections, gathering information from distributed brain networks as the GNW theory has postulated, but may be less responsible for information differentiation in terms of spatiotemporal variability in brain modules, failing to provide strong support for the IIT. Together, our study provides an analytic framework based on graph-theoretical tools to locate the NCC architecture, which complements our understanding about the anatomical footprints underlying consciousness by demonstrating their superior ability not only in functional integration, but also in dynamic functional diversification with modular processors. Furthermore, our results showed that the NCC candidates are not limited within the posterior cortical regions, but are distributed in wide-spread frontal-parietal cortices, which contribute to the ongoing discussion of whether the NCC are residing in the anterior or posterior regions^37,38^.

Interestingly, we demonstrated that the identified NCC members are associated with regional expression of genes that are enriched in neuronal-related pathways predominantly involved in synaptic functions and transmission, which is in line with the previous findings reporting a strong link between transcription profiles and macroscale brain topology ^39-41^. The spatial layout ^42^, connectivity ^41^, as well as the information synergy gradient ^43^ of the association networks, such as the DMN and FPN, have been observed to be captured by coupled transcription profiles of genes enriched in synapse-related functions. These findings highlight the critical role of synaptic functions in information communications across the brain networks, which may also provide a potential neurobiological underpinning of the NCC’s involvement in higher-order cognitive functions, as revealed in our meta-analysis.

Our observations from dynamic analysis provide further evidence supporting the functional implications of the NCC architecture in the temporal domain. Based on theoretical and simulation evidence, the GNW theory predicts two dynamic states or stages for a stimulus to access consciousness ^6^: an “ignited” conscious state when the input signal is strong enough to ignite a sustained, metastable distributed neural representation of the current conscious contents; and a subliminal processing state during which the incoming activity is propagated through primary sensory areas inducing only a progressively decaying activity in higher-order regions. Intriguingly, this fits well with the roles that our findings assigned to the two brain states identified herein. Specifically, we demonstrated that within the brain state of higher temporal stability of modular organization, a significant percentage of the most stable regions coincides with the identified NCC. In contrast, the other brain state showing significantly lower stability is mainly steered by primary sensory areas. Furthermore, our results pinpoint the default midline areas to be the temporally most stable sites during the temporally more stable state, which refines our understanding of the critical value of DMN areas in maintaining neural representations that access consciousness. This observation is also consistent with recent studies showing that the default mode regions that are distant from sensory input have the longest timescale to accumulate and process information, which may facilitate functional integration over a slow timescale ^44-46^.

In supportive of NCC’s roles in human consciousness, we demonstrated their selective vulnerability during anesthesia-induced loss of consciousness. During loss of consciousness, although static FC strength of both NCC and local connections decreased, dynamic analysis revealed that connectivity within the NCC was particularly impacted during the temporally more stable state, which is consistent with previous dynamic connectivity studies in patients under pharmacological or pathological unconsciousness, showing more extensively disruptions in states that potentially relevant for conscious processing ^47-49^. Furthermore, we also found that the identified NCC members exhibited different anesthesia-related changing patterns in terms of the two network measures, FCS and MV. Specifically, FCS reductions were specific to the DMN midline areas, while decreases in MV were observed predominately in prefrontal cortices, suggesting that these two sets of regions within the NCC may play complementary roles in supporting consciousness. This observation is in line with previous hypothesis proposing that the areas constituting the neural substrates supporting conscious are “neither identical nor redundant” ^8^, and offers novel insight into the understanding of functional specificity of NCC members. The FCS indicates a region’s ability in integrating or amplifying segregated information, while the temporal modular variability reflects its temporal flexibility in selecting or broadcasting information with a wide range of local processors. Thus, the DMN midline regions may be well-suited for functional integration rather than functional diversification, while the prefrontal areas may function in a distinct way in supportive of conscious processing. Indeed, recent insights from connectivity decomposition analysis found that the DMN is situated at the top of the topographical hierarchy in brain connectivity space, supporting its essential role acting as cortical hubs with a maximal distance from primary systems to integrate and represent multi-dimensional information ^50^. In contrast, the fronto-parietal system was found to occupy a position sitting between the unimodal and default mode networks, suggesting it may function as a connector to collect bottom-up information and distribute top-down information ^50^. Notably, the medial prefrontal region exhibited reduction in both FCS and temporal MV associated with propofol anesthesia, indicating a crucial role of this anterior cluster in contributing to conscious experience. Taking together, these observations provide evidence for the pivotal contribution of the identified putative NCC, which covered most key structures predicted by both GNW and IIT, in supporting human consciousness, and more importantly, highlight the potential differential roles between prefrontal GNW regions and posterior IIT regions during propofol-induced loss of consciousness ^37^.

There are a few limitations to note in our study. First, given the relatively limited scan duration of our primary dataset (8 mins), NCC identification was only performed on group-level, which would underestimate potential individual differences in the topography and topology of the NCC architecture. To mitigate this concern, we conducted individualized analysis on the MSC dataset and validated that the individual-level NCC distribution was similar with that on group-level. Nevertheless, future research would benefit to consider constructing individualized NCC to predict between-subject variability in conscious access and related cognitive abilities. Second, we used a single type of anesthetic (a GABA_A_ receptor agonist) to induce loss of consciousness. It is unknown whether our observation of the disruptions during propofol-induced anesthesia would generate to other types of anesthetics, or other forms of unconsciousness. Previous studies have consistently related changes in the prefrontal and default mode regions, which are key members of the NCC identified herein, to unconsciousness induced by various types of anesthetic or disorders of consciousness ^13,17,51,52^, attesting to the generality of NCC’s selective vulnerability. However, NCC members could still exhibit different spatiotemporal changing patterns during different forms of unconsciousness, which warrants future investigations.

## Material and methods

### Anesthesia dataset

Twenty-one patients (male/female: 9/12; age: 32–64 years) who were selected for elective resection of pituitary microadenoma via a transsphenoidal approach (<10 mm in diameter without sella expansion by radiological and plasma endocrine indicators) attended the experiments. The participants had American Society of Anesthesiologists physical status grades of I or II and no history of craniotomy, cerebral neuropathy, vital organ dysfunction, or administration of neuropsychiatric drugs. The procedure was approved by the institutional review board of Huashan Hospital, Fudan University. All participating subjects provided written informed consent.

### Anesthesia protocol

The participants were administered intravenous propofol anesthesia. By target-controlled infusion, we sustained 1.3 μg/mL effect-site concentration for a Ramsay sedation scale of 3–4. The patients were then administered remifentanil (1.0 μg/kg) and succinylcholine (1.5 mg/kg) to facilitate endotracheal intubation. For general anesthesia, we maintained a stable effect-site concentration (4.0 μg/mL) that reliably kept patients unconscious. During general anesthesia, intermittent positive pressure ventilation was used with tidal volumes of 8–10 mL/kg, respiratory rate of 10–12 beats per minute, and PetCO_2_ of 35–40 mmHg. In the postoperative follow-up, no subject reported intraoperative awareness during the fMRI scanning and surgical procedure.

### Image acquisition

Participants were scanned on a Siemens 3T scanner (Siemens MAGNETOM, Germany) to acquire high-resolution T1-weighted anatomical images (echo time (TE) = 5 ms, repetition time (TR) = 1,000 ms, slice thickness = 1.0. mm, 176 slices, image size = 448×512, FOV = 219×250 mm^2^, flip angle = 90°) and resting-state fMRI images using gradient-echo echo-planar imaging (EPI) sequence (TE = 30ms, TR = 2,000ms, slice thickness = 5.0mm, number of slices = 33, repetition = 240, image size = 64×64, FOV = 210×210mm^2^, flip angle = 90°). Resting-state fMRI images were scanned eyes-closed for 8 minutes during wakefulness, sedation, and general anesthesia, respectively. Before sedation scanning, the subjects were administered 500 mL intravenous hydroxyethyl starch to prevent hypotension caused by propofol-induced vasodilation.

### Validation dataset

A secondary, independent dataset including fMRI images from ten subjects was downloaded from Midnight Scan Club website (MSC, https://legacy.openfmri.org/dataset/ds000224/) for validations. Briefly, the participants were scanned on a Siemens 3.0T Tim Trio (software version syngo MR B17) to acquire high-resolution T1-weighted anatomical images (TE = 3.7 ms, TR = 2,400 ms, slice thickness = 0.801 mm, 256 slices, image size = 224×256, FOV = 179×205 mm^2^, flip angle = 8°) and resting-state fMRI images using gradient-echo echo-planar imaging (EPI) sequence (TE = 27 ms,TR = 2,200 ms,slice thickness = 4.0 mm,number of slices = 36,repetition = 818, image size = 64×64,FOV = 256×256 mm^2^ flip angle = 90°). Resting-state fMRI images were scanned eyes-open for 30 minutes per session, totaling 300 min per subject. Details of the MSC dataset are published elsewhere ^53^.

### fMRI image processing

Resting-state fMRI images from the anesthesia dataset were preprocessed using AFNI ^54^. After discarding the first five frames, fMRI images were corrected for different slice timing, realigned, coregistered with the high-resolution anatomical images, and spatially normalized to the standard template brain (Talaraich stereotactic space). Head motion and signals from the white matter and cerebrospinal fluid were regressed out to control for physiological and non-neural noise. Band-pass filtering (0.01–0.1 Hz) and spatial smoothing (6 mm FWHM) were then applied. Since the human subjects underwent fMRI scans from wakefulness to anesthesia, the head motions tend to be larger while awake; to mitigate motion-related confounding, subjects with extra head motions were discarded from further analyses ^55,56^. The exclusion criteria included: (1) averaged head translation/rotation >1.5 mm/°; (2) 4-min longer time points with frame-wise displacement (FD) >0.4 mm. We also confirmed that there were no significant differences in the numbers of time points with excessive FD (*F*(2, 24) = 0.99, *P* = 0.38) and averaged head motion (*F*(2, 24) = 1.86, *P* = 0.17) among doses.

For the MSC validation dataset, preprocessed fMRI images were downloaded directly from MSC website. Note that one subject is reported repeatedly falling asleep and exhibited several non-negligible eye closures, along with increasing head motion ^53^, thus was excluded, remaining nine subjects for further analyses.

### Static functional network construction

To construct functional brain networks, we first parcellated the brain according to a predefined brain atlas, which yielded 246 regions of interest the human brainnetome atlas, ^57^. For each subject, we averaged the voxel-wise time course within each region and computed the Pearson correlation to obtain a functional connectivity (FC) matrix. Density thresholds of S = 15% was applied to remove as many spurious correlations as possible while maintaining fully connected brain networks.

### Dynamic functional network construction

Dynamic FC was estimated using a sliding window approach. We used a window length of 25 TRs slides in steps of 1 TR individually, resulting in 211 time windows (TWs) at each of the three conscious levels for the propofol anesthesia dataset and 8156 TWs for the validation dataset. We then calculated the Pearson’s correlation coefficients for each pair of regions from the windowed time course segments. The resulting windowed correlation matrices were thresholded using connectivity densities of 15% to generate dynamic functional networks.

### Rich club identification

The rich-club organization represent a core set of highly-connected regions that tends to be more densely interconnected in a network. To detect rich-club organization in a weighted network, we first computed the FC strength of each brain region as the average of FC with all the other regions, and calculated the rich-club coefficient as follow ^58,59^ :

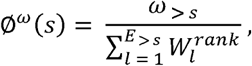

Where *ω* _> *s*_ is the sum of connectivity strength between nodes with nodal strength higher than s, *E* _> *s*_ is the number of these connectivity, and 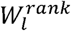 is the sum of the top *E* _> *s*_ strongest connectivity across the network. By dividing Ø ^*ω*^ (*s*) by 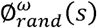, a normalized 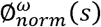 is generated, where 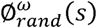 is the averaged rich-club coefficient of 1000 random networks with the same nodal degree and strength distribution. A network appears to have the rich-club organization if 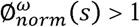 for a continuous range of s. Permutation test was performed to assign a P value at each s by comparing the observed Ø ^*ω*^ (*s*) with the null distribution of 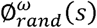 obtained from random networks. Bonferroni method was used to correct multiple comparisons.

### Modular structure detection

We assessed the modular structure by applying modularity analysis on static or dynamic brain networks ^22,60,61^. To achieve an optimal module partition, we applied a two-step procedure similar to that described by Rubinov and Sporns ^62^. Briefly, the modular partition was first estimated using the Louvain algorithm 100 times, followed by a fine-tuning algorithm that was performed repeatedly until the modularity of the partition no longer increased; a consensus partition was then identified with the highest modularity. To choose an appropriate value for the resolution γ parameter in the modularity analysis, we repeated the above two-step procedure across a range of γ values (1–2 in steps of 0.1); for modular partition obtained for each γ value, we computed the variation of information (VI) with those identified at the neighboring γ values, and γ parameter showing the lowest VI was selected as it provides the most robust estimates of topology across these iterations ^22,26^.

### Modular variability

To estimate the variability of spatial affiliations between different modular partitions, for a given brain region *k*, we calculated the modular variability (MV) between modular affiliation *i* and *j* obtained in different subjects or time windows as follows^25,63^:

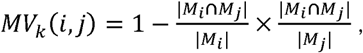

where *M*_*i*_ and *M*_*j*_ denote the module label to which region *k* belongs in modular partitions *i* and *j*, respectively. *M*_*i*_ ∩ *M*_*j*_ represents the mutual region set between modules *M*_*i*_ and *M*_*j*_, and |*M*_*i*_ ∩ *M*_*j*_ | denotes the number of regions in the common region set. |*M*_*i*_ | and |*M*_*j*_| denote the number of regions in modules *M*_*i*_ and *M*_*j*_, respectively. The average modular variability for node *k* across all *n* modular partitions can then be calculated as:

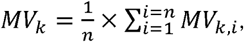

where 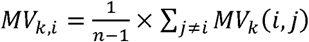 denotes the modular variability for region *k* between modular partition *i* and all other partitions.

### Dynamic state detection

Recent evidence suggested that fluctuations of neuronal activity could be characterized by highly structured connectivity patterns that reoccur over time, which have been described as “brain states” ^64-67^. Since the present study focused on spatial variability of modular organization, we developed a new method to detect brain states at individual and group levels based on the temporal variability of modular structure in functional brain networks.

To identify dynamic states in each individual during different levels of responsiveness, we first detected the modular structures of each time-varying brain network using the Louvain modularity algorithm in GRETNA toolbox ^68^. To estimate the similarity of modular structures between different TWs, for each brain region *k*, we generated a similarity matrix, iMV(w × w), by inversing the across-time *MV*_*k*_(*i, j*) values (w: number of time windows). Modularity analysis was then applied to this similarity matrix, iMV(w × w), to identify modules that might correspond to sets of TWs with similar brain modular organization. The first-level analysis identified a range of 2 to 4 states for each region in more than 90% of subjects.

To reveal brain states that reoccurred across subjects and different levels of responsiveness, a second-level analysis was performed based on the individual-level brain states. First, for each subject and each condition, we averaged the inversed across-time *MV*_*k*_ of each region across TWs within each first-level brain state. The TWs of the largest averaged inversed across-time *MV*_*k*_ were labeled as 1, while the remaining TWs were labeled as 0. We then concatenated the TWs across all subjects and all conditions to generate a matrix, *iMV*_*th*_ (n × r), where n is the number of TWs multiplied by the number of subjects and conditions and r is the number of brain regions. Each column in the *iMV*_*th*_ (n × r) matrix represented whether a given brain region was temporally stable in its module in current TW. Finally, a similarity matrix, S(n × n), was computed as the inversed Euclidean distance between each pair of rows of *iMV*_*th*_ (n × r). We then performed a modularity analysis on S(n × n) to detect group-level brain states. Within each state, temporal stability was estimated as the proportion of 1s in the matrix *iMV*_*th*_ (n × r).

### Term-based meta-analysis of the NCC structure

Meta-analysis was conducted to decode the potential functionality of the identified NCC structure using NeuroQuery ^31^ and NeuroSynth ^32; www.neurosynth.org^ database. The NeuroQuery term database was first manually filtered to exclude terms related to anatomical structure, disorders and some acronyms and serial number, then encoded to obtain each term’s activation map by integrating findings of existing term-related articles and calculating the number of related articles (denoted as frequency) as weights in the map. Similarity was then calculated between the NCC map and each activation map, measuring the spatial pattern similarity between them and the term’s frequency, after which all terms were sorted in a descending order by their similarities. This procedure was conducted according to NeuroQuery’s pipeline released on github (https://github.com/neuroquery). 50 terms with top similarity were displayed in a word cloud with similarity as weights. The identified NCC map was also decoded using the decoder tool provided in NeuroSynth (https://neurosynth.org/decode/), generating terms in descending order of similarity. After that, 50 terms with top similarity were displayed in a word cloud after manually discarding terms related to anatomical structures and disorders.

### Transcriptional genetic basis identification of NCC

Leveraging the unprecedented regional gene expression measurements of six neuropsychiatric and neurological healthy adult human brains (24-57 years), which was contributed by Allen Brain Institute (http://human.brain-map.org/; RRID: SCR_007416) ^1,33^, we further explore the transcriptional basis underlying the NCC structure. Since only 2 donors have data for the right hemisphere, we used data from the left hemisphere for reliability consideration, in consistent with prior studies ^1^. A recently published practical guide for processing Allen Human Brain Atlas (AHBA) dataset was applied ^69^. Specifically, we first re-annotated probes to genes using the latest database in Re-annotator; removed expression values below background. For genes with multiple probes, we take differential stability measurement to estimate the expression value. Samples from AHBA were mapped to Brainnetome Atlas regions for left cortex and left subcortex separately. Scaled robust sigmoid (SRS) normalization were used to account for inter-subject differences and outliers. Differential stability was used to select genes consistently expressed across six brains. Finally, a data matrix of 123 regions x15655 genes was generated.

Differential expression between NCC and non-NCC region were then assessed using the quantile-normalized gene expression matrix across 123 left-hemispheric cortical and subcortical regions, results in a ranked gene list that measures the extent to which the genes were differential expressed. The Limma R package was utilized here and no significance threshold was set ^70^.

Functional annotation for the whole ranked gene list generated above was assessed using Gene ontology (GO) biological pathway and KEGG pathway enrichment analysis, which is implemented with GAGE package in R version 4.0. Neuronal-related pathways were shown in a word cloud at the significance of *P* < 0.05.

### Statistical Analysis

To detect differences in temporal stability between the two dominant states, a paired t test was performed (*P* < 0.05). To evaluate spatial overlap between stable regions of each state and NCC areas, a nonparametric permutation test was used, whereby the ratio of NCC members overlapping with stable regions (> mean) within each state was calculated. A total of 1,000 random permutations were generated independently; for each permutation, stability values were randomly permuted across brain regions within each state, and the test statistic of interest was recalculated, generating a distribution of 1,000 values from the permuted data. The *P* value was determined by comparing the observed value with the permutation-generated distribution. *P* values of less than 0.05 was considered statistically significant. To test which state involves more NCC members, we calculated between-states difference in NCC overlapping ratio, and performed the non-parametric permuation test using the same procedure as above-mentioned.

To evaluate the effect of propofol-induced anesthesia, a one-way, repeated-measures ANOVA, with conscious condition (awake, sedation and anesthesia) as the within-subject factor, was applied to mearsures of connectivity strength (NCC, feeder and local connections) and modular variability (NCC and non-NCC members), respectively, followed by independent, post hoc, paired t tests between every pair of conditions. Significance was considered at *P* < 0.05 and corrected for multiple comparisons using Bonferroni correction. Furthermore, network-based statistics (NBS) analysis was implemented using GRETNA MATLAB Toolbox ^68^ between pairs of conscious conditions, to reveal whole-brain network-based features of loss of consciousness with significance set at *P* < 0.005. Regional-wise comparisons of across-time MV between every pair of conditions was performed by paired t tests with significance set at *P* < 0.05 and corrected using False Discovery Rate (FDR) correction. The NBS and regional-wise results were presented using BrainNet Viewer ^71^.

## Supplemental figure legends

**Figure S1.**
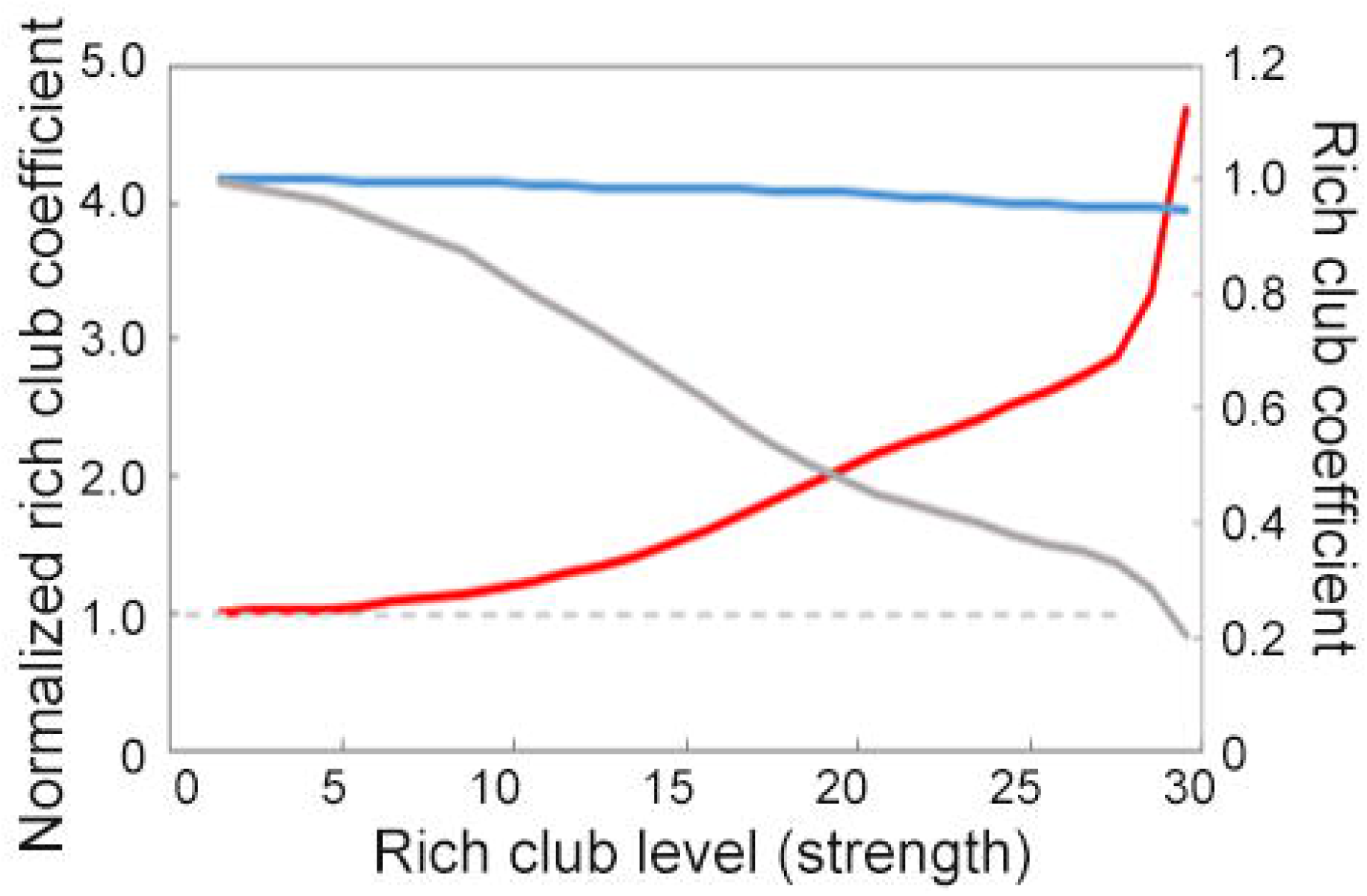
Rich-club coefficient curves for functional brain networks during awake consciousness (blue line), and for random networks (gray line) and the normalized rich-club coefficients (red line).

**Figure S2.**
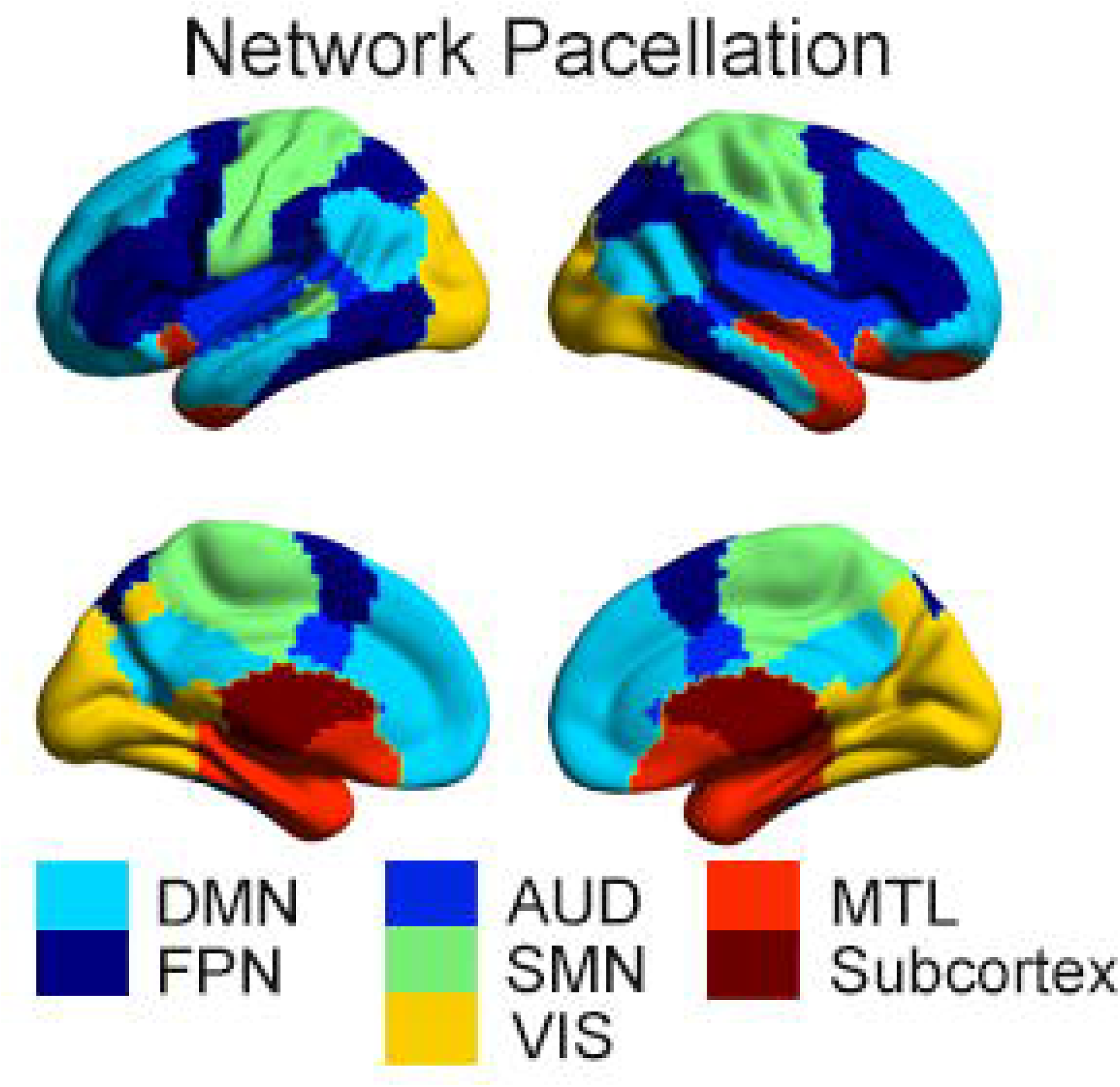
Spatial distributions of seven identified modules of the group-averaged functional brain networks during wakefulness. *DMN: default mode network, FPN: frontoparietal network, AUD: auditory network, SMN: sensorimotor network, VIS: visual network, MTL: medial temporal lobe, Subcortex: subcortical network*.

**Figure S3.**
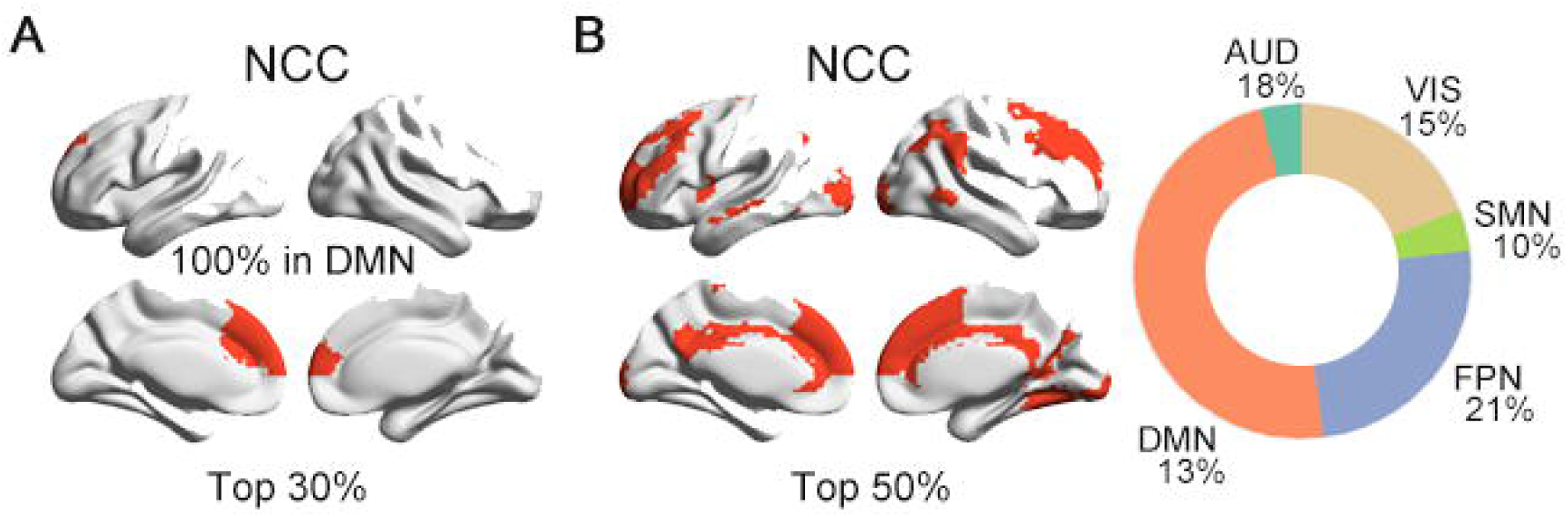
NCC templates identified by overlapping maps of the three topological metrics using different percentile (top 30% and top 50%) of regions with top rich-clubness and MV.

**Figure S4.**
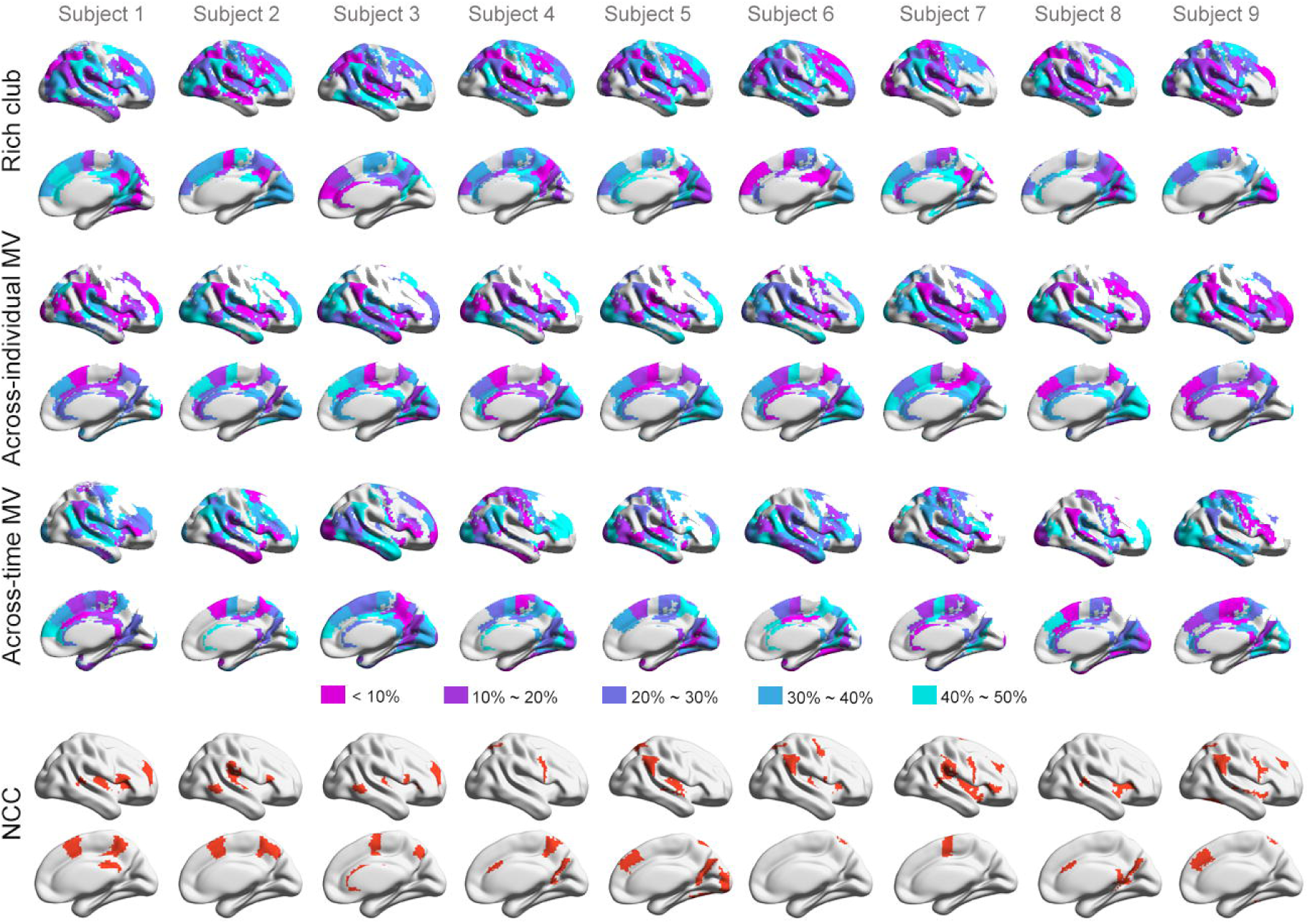
Individual-level maps of rich-club, across-time MV, across-individual MV, and identified NCC in MSC dataset.

**Figure S5.**
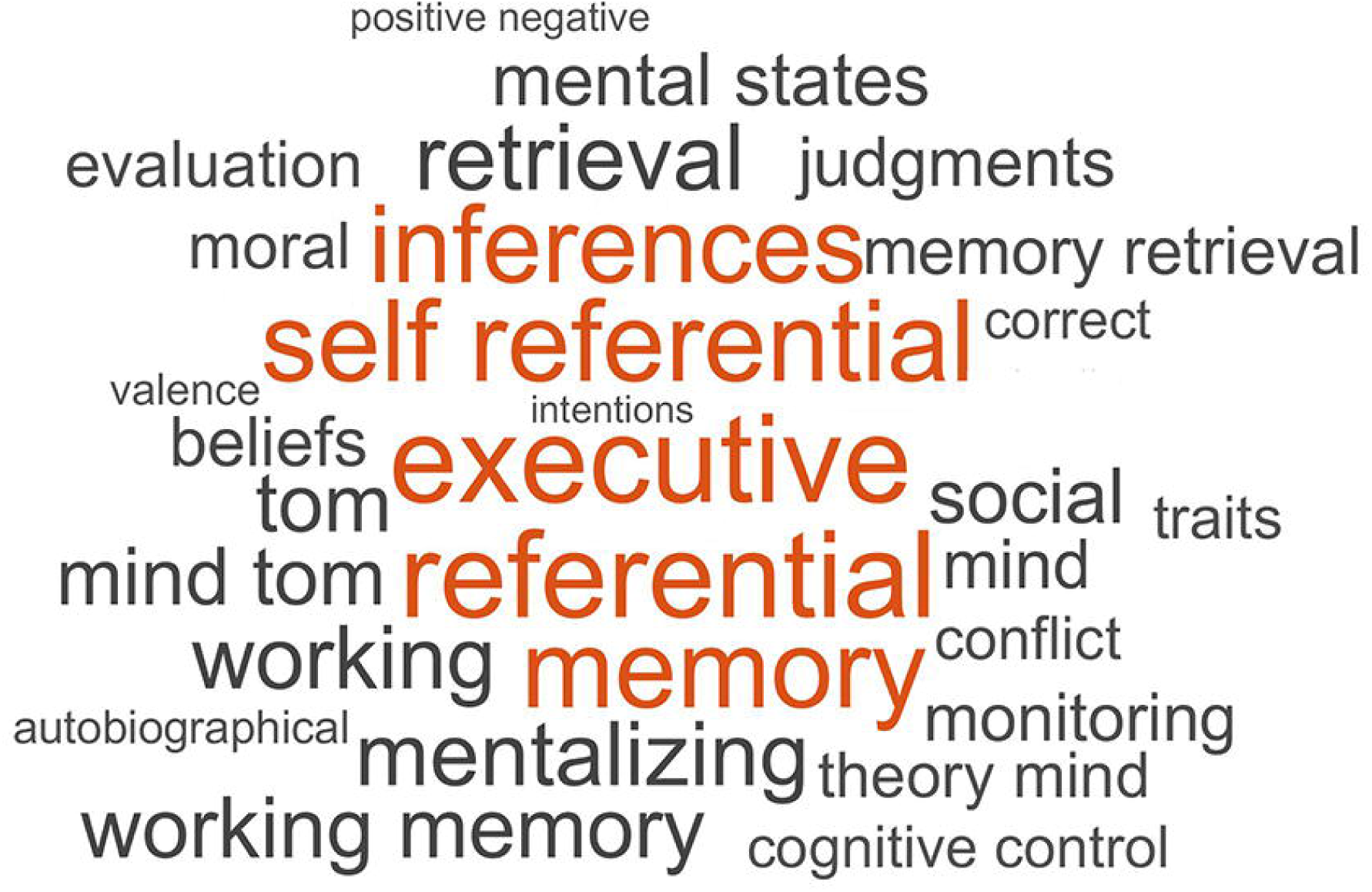
Meta-analysis results based on the NeuroSynth database.

**Figure S6.**
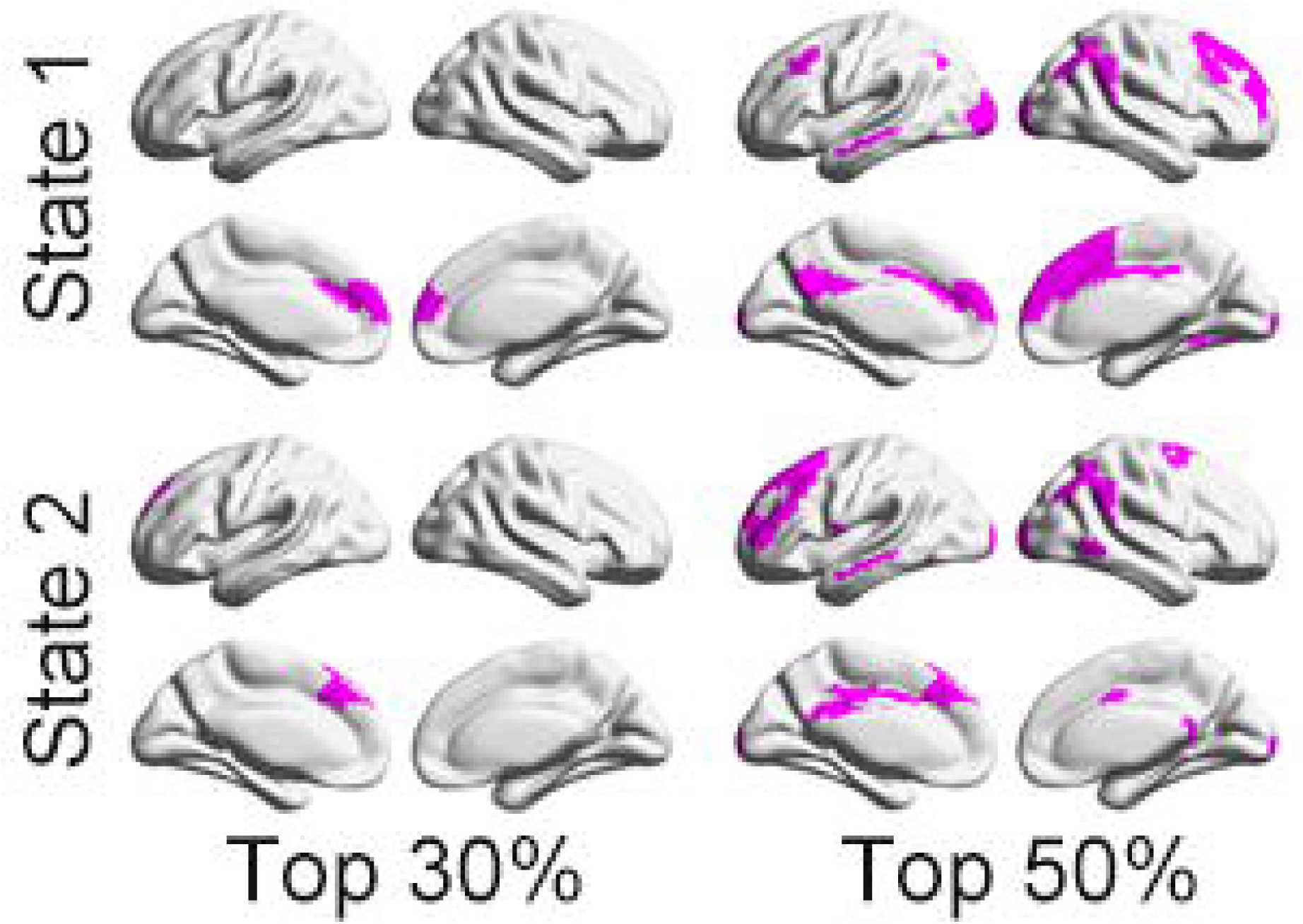
Overlaps between the identified NCC space of different percentile (top 30% and top 50%) with stable regions in each dynamic state.

**Figure S7.**
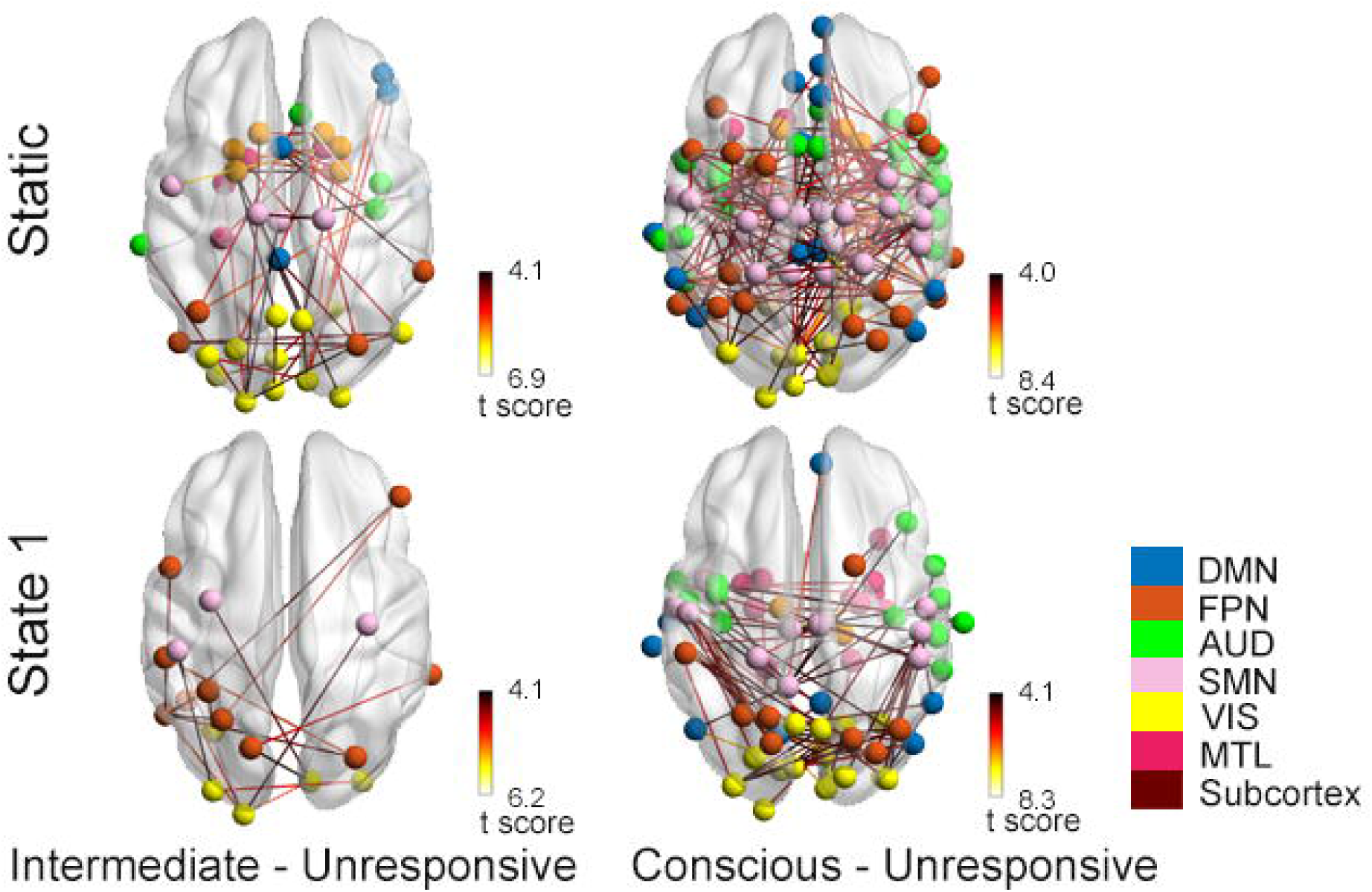
Network-based statistics (NBS) revealed significant reductions in local connections between different conscious conditions during static (upper panel) and dynamic state 1 (bottom panel).

**Figure S8.**
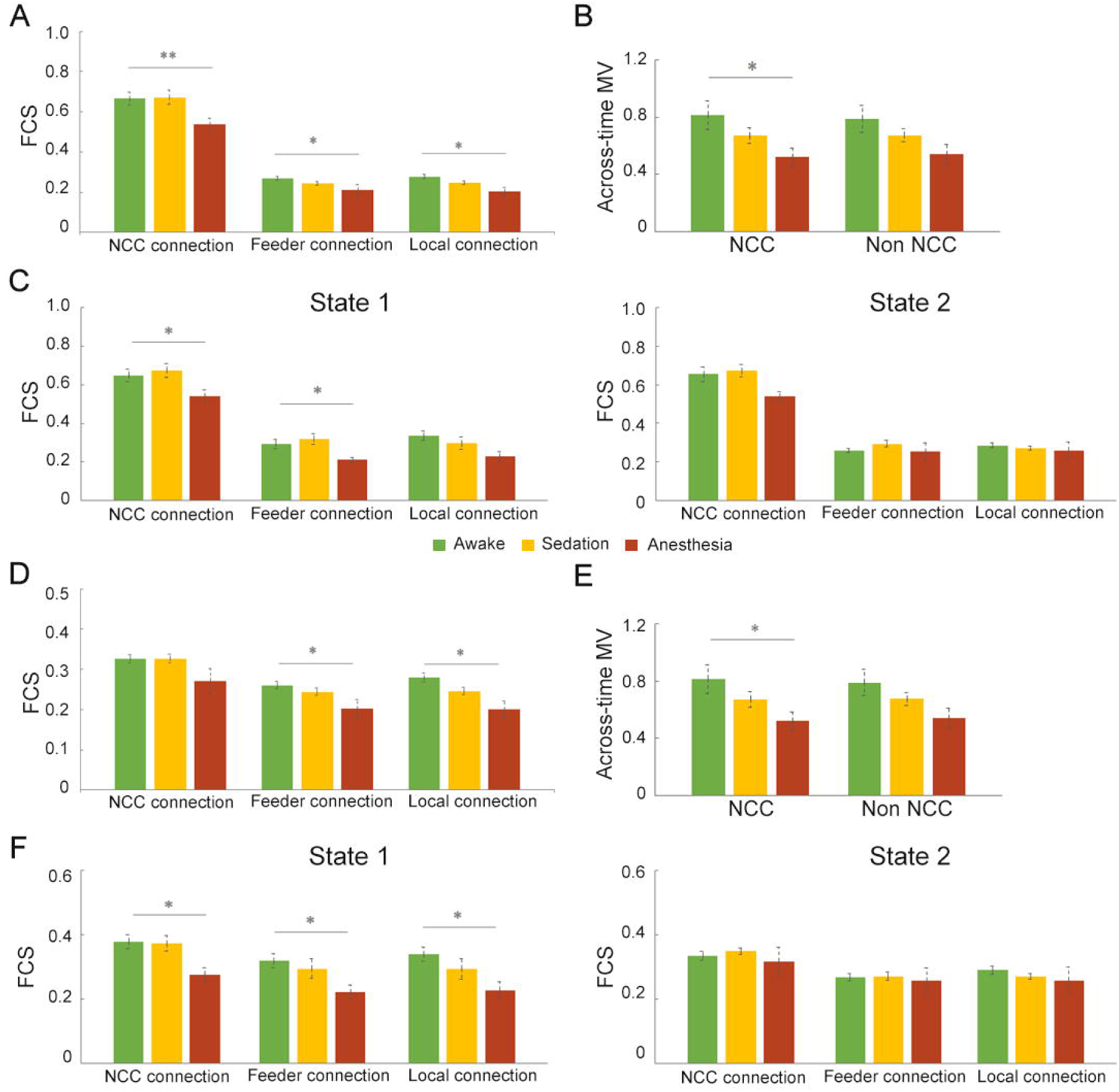
NCC signatures track loss of consciousness during Propofol anesthesia using two different percentiles: top 30% (A-C) and top 50% (D-F). *\*, Puncorrected < 0*.*05; **, Pcorrected < 0*.*05*.

## Funding Statement

This work was supported by the National Natural Science Foundation of China (https://www.nsfc.gov.cn/, grant numbers 82072000 and 81671769 to [XL]); the Central University Basic Research Fund of China (https://www.edu.cn/, grant number HIT. NSRIF. 2020042 to [XL]); the Natural Science Foundation of Heilongjiang Province, China (http://kjt.hlj.gov.cn/, grant number LH2019H001 to [XL]); and the Medical Guidance Supporting Project (http://stcsm.sh.gov.cn/, grant numbers 17411961400 and 20Y11906200 to [JZ]) from the Municipal Science and Technology Committee, Shanghai, China.

## Conflicts of Interest

The authors have declared that no competing interests exist.

## Data availability statement

The data under propofol administration that support the findings of this study are not publicly available due to restrictions imposed by the administering institution and privacy of the participants. The authors will share them by request from any qualified investigator after completion of a data sharing agreement. The MSC data is a publically available dataset at Midnight Scan Club website (https://legacy.openfmri.org/dataset/ds000224/).

## Code availability statement

Code for data cleaning and analysis will be updated as part of the replication package once the paper has been conditionally accepted.

## Notes

https://legacy.openfmri.org/dataset/ds000224/

### Competing Interest Statement

The authors have declared no competing interest.

